# Metabolomic profiling of mouse mammary tumor derived cell lines reveals targeted therapy options for cancer subtypes

**DOI:** 10.1101/796573

**Authors:** Martin P. Ogrodzinski, Sophia Y. Lunt

**Affiliations:** Department of Biochemistry and Molecular Biology, Michigan State University, East Lansing, MI, USA; Department of Physiology, Michigan State University, East Lansing, MI, USA; Department of Chemical Engineering and Materials Science, Michigan State University, East Lansing, MI, USA

## Abstract

Breast cancer is a heterogeneous disease with several subtypes that currently do not have targeted therapy options. Metabolomics has the potential to uncover novel targeted treatment strategies by identifying metabolic pathways required for cancer cells to survive and proliferate. Here, we used tumor-derived cell lines derived from the MMTV-Myc mouse model to investigate metabolic pathways that are differentially utilized between two subtypes of breast cancer. Using mass spectrometry-based metabolomics techniques, we identified differences in glycolysis, the tricarboxylic acid cycle, glutathione metabolism, and nucleotide metabolism between subtypes. We further show the feasibility of targeting these pathways in a subtype-specific manner using metabolism-targeting compounds.

## INTRODUCTION

Breast cancer is a heterogeneous disease with subtypes that vary by morphology, receptor status, and gene expression profiles.^1,2^ This diversity impacts treatment, as one therapeutic strategy will not work for all patients. Targeted therapies include endocrine therapy for patients with estrogen receptor positive (ER+) breast cancer, as well as monoclonal antibodies/inhibitors against human epidermal growth factor receptor 2 (HER2) for HER2+ breast cancer.^3^ Unfortunately, targeted therapies are not available for every breast cancer subtype, and drug resistance and relapse remain problematic.^4,5^ Therefore, it is critical to identify additional therapeutic targets for all subtypes of breast cancer, and investigating cancer metabolism has the potential to meet this need.^6^

Cancer cells exhibit metabolic differences compared to normal cells, and dysregulated metabolism is considered to be a hallmark of cancer.^7^ Central carbon metabolism, which includes pathways such as glycolysis, the tricarboxylic acid (TCA) cycle, the pentose phosphate pathway (PPP), and amino acid metabolism, is dysregulated in cancer cells and fuels survival and proliferation.^8^ Previous work has shown that metabolic dysregulation can be specific to different subtypes of breast cancer. For example, HER2+ and triple negative breast cancer (TNBC) have been shown to upregulate glutaminolysis compared to ER+ breast cancer,^9,10^ and TNBC cell lines and xenograft models are sensitive to glutaminase inhibition.^11^,^12^ Differential utilization of metabolic pathways between subtypes of cancer therefore represent potential targets that can be leveraged to develop novel treatment strategies.

The diversity observed in human breast cancer can be modeled by the MMTV-Myc mouse model.^13^ MMTV-Myc mice develop mammary tumors that display heterogeneity in both histology and gene expression.^14^ Histological subtypes of the MMTV-Myc model have previously been correlated with human subtypes based on global gene expression.^15^ For example, MMTV-Myc epithelial-mesenchymal-transition (EMT) tumors correspond to the claudin-low subtype of human breast cancer, a subtype which currently has no targeted therapy options and is generally associated with a poor prognosis.^16^ Additionally, compared to MMTV-Myc EMT tumors, MMTV-Myc papillary tumors display increased Myc signaling pathway activity,^14^ which is also amplified in 15.7% of human breast cancers and is more common in high grade tumors and the basal-like subtype.^17^ As a transcription factor, Myc affects numerous biological processes including metabolism.^18,19^ Notably, Myc expression regulates several genes in glucose, amino acid, and nucleotide metabolism.^20–22^ Therefore, investigating metabolism of the MMTV-Myc model system may reveal metabolic features common to human cancer and could present new targeted therapeutic options.

Here, we present a study investigating the metabolic profiles of two histologically distinct breast cancer subtypes, EMT and papillary, from the MMTV-Myc mouse model. We developed a workflow to investigate and target the metabolic profiles of mouse mammary tumor subtypes (**Figure 1**). Cell lines were derived from primary EMT and papillary tumors, and polar metabolites were extracted and analyzed using an optimized liquid chromatography tandem mass spectrometry (LC-MS/MS) method to measure a wide range of metabolites.^23^ Based on metabolic profiles, drugs most likely to be effective against each subtype were selected and tested. We found that, compared to the papillary subtype, the EMT subtype demonstrated increased glutathione and TCA cycle metabolism, while the papillary subtype had increased nucleotide biosynthesis compared to the EMT subtype. Targeting these distinct metabolic pathways effectively inhibited cancer cell proliferation in a subtype-specific manner. These results demonstrate the potential utility of metabolic profiling to develop novel personalized therapeutic strategies for different subtypes of breast cancer.

**Figure 1:**
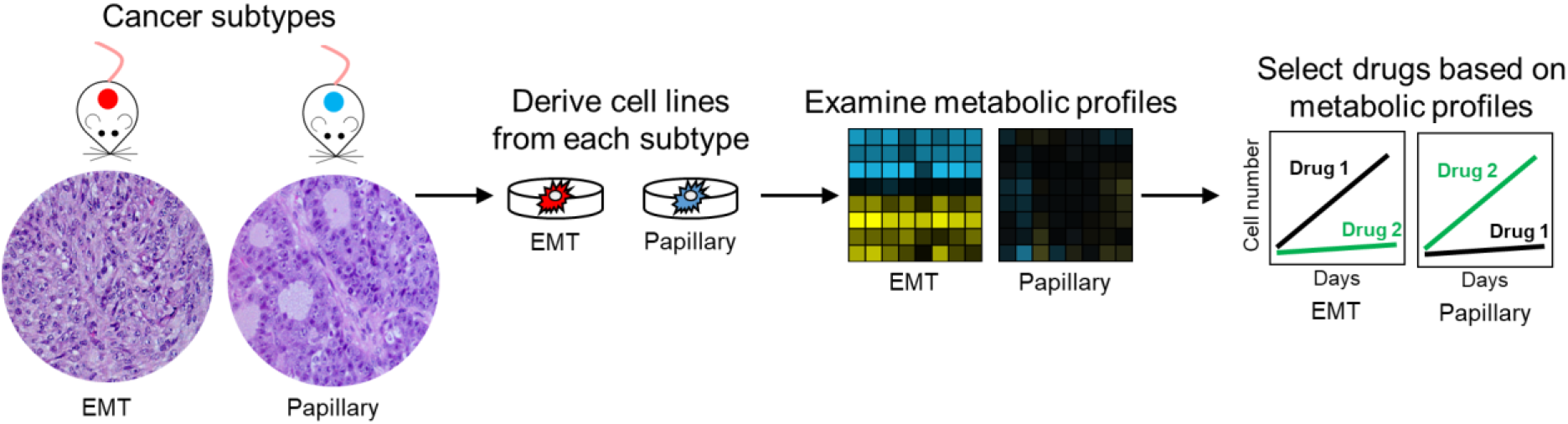
Schematic overview of the experimental design for drug selection based on breast cancer subtype-specific metabolism. The epithelial mesenchymal transition (EMT) and papillary tumors are histologically distinct mouse mammary tumor subtypes from the MMTV-Myc mouse model. Cell lines derived from tumors can be used to determine metabolic pathways that can be used to select drug candidates for each subtype.

## RESULTS

### Relative metabolite levels between histologically distinct subtypes of MMTV-Myc mouse mammary tumors define metabolic pathways of interest

To determine metabolic profiles of histologically distinct mouse mammary tumor subtypes, polar metabolites were extracted from tumor-derived cell lines and quantitated using LC-MS/MS. We found metabolites involved in several central carbon metabolic pathways to be differentially abundant between EMT and papillary tumor-derived cell lines (**Figure 2**).

**Figure 2:**
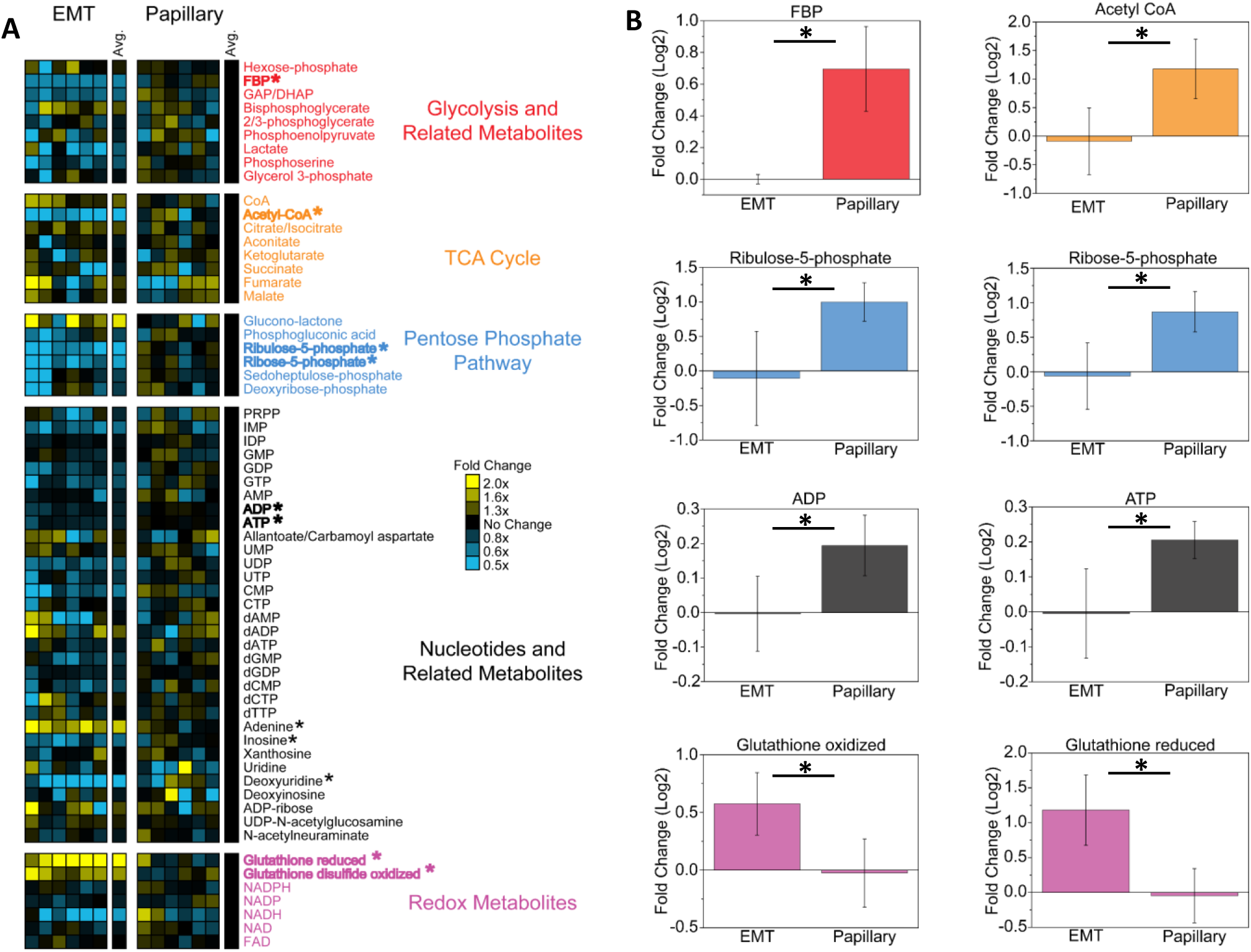
Metabolite pool sizes are different between EMT and papillary tumor derived cell lines. (**A**) Heatmap indicating relative metabolite differences between EMT and papillary tumor derived cell lines. Metabolites are sorted by relationship to metabolic pathways. Yellow and blue boxes indicate increased or decreased metabolite levels relative to the average of the papillary subtype, respectively. Data for each sample is normalized to the average signal for all metabolites in the analysis. Metabolites with statistically significant differences (p-value < 0.05) are bolded and marked with asterisks (*). (**B**) Representative bar graphs of metabolites with statistically significant differences between EMT and papillary subtypes. Data are displayed as means ± S.D., N = 6.

In the EMT subtype, both oxidized and reduced forms of glutathione, a key metabolite in redox homeostasis, are elevated (**Figure 2B**). Increased levels of both reduced and oxidized glutathione imply that the EMT subtype has elevated glutathione biosynthetic activity. This could reflect a greater dependency on glutathione biosynthesis in the EMT cells and targeting glutathione biosynthesis would therefore be more effective against the EMT subtype. Metabolites increased in the papillary subtype include fructose bisphosphate (FBP; glycolysis); acetyl-CoA (TCA cycle); ribulose-5-phopsphate and ribose-5-phosphate (PPP); and adenosine diphosphate (ADP) and adenosine triphosphate (ATP; nucleotide metabolism; **Figure 2B**). Additional analysis using isotope labeling is required to determine whether metabolites are present at higher levels due to higher production or lower consumption.

### Isotope labeling through the TCA cycle is increased in the EMT subtype

Elevated acetyl-CoA and FBP levels in the papillary subtype (**Figure 2B**) could be explained by either increased glycolytic activity or decreased TCA cycle activity, since these pathways contribute to the production or consumption of these metabolites, respectively. To further investigate these relative metabolic pathway activities, we performed stable isotope labeling using ^13^C-glucose and ^13^C-glutamine. Isotope labeling studies show the rate at which these metabolites are incorporated into different metabolic pathways, enabling comparison of relative metabolic pathway activities between samples. Isotope labeling patterns complement metabolic pool size measurements to reveal more complete metabolic profiles.^24^ We find the papillary subtype has proportionally lower abundance of ^13^C-labeled glycolysis and TCA cycle intermediates from ^13^C-glucose compared to the EMT subtype (**Figure 3; Supplementary Table 1**). Therefore, the increased abundance of FBP and acetyl-CoA in the papillary cells are likely due to decreased TCA cycle activity in this subtype compared to the EMT cells. Notably, the labeled fraction of 2/3 phosphoglycerate (66% in EMT vs. 56% in papillary), alpha-ketoglutarate (40% in EMT vs. 28% in papillary), succinate (73% in EMT vs. 42% in papillary), fumarate (54% in EMT vs. 33% in papillary), and malate (44% in EMT vs. 32% in papillary) are each higher in the EMT cells when ^13^C-glucose is used as the labeled carbon source (**Supplementary Table 1**). When ^13^C-glutamine is used as the labeled carbon source, most TCA cycle metabolites do not demonstrate a significant difference in labeling between the EMT and papillary cells (**Supplementary Figure 1; Supplementary Table 2**). These results indicate that EMT cells increase glucose flux through the TCA cycle to a greater degree than papillary cells. Thus, targeting the TCA cycle is likely to be more effective in the EMT subtype compared to the papillary subtype.

**Figure 3:**
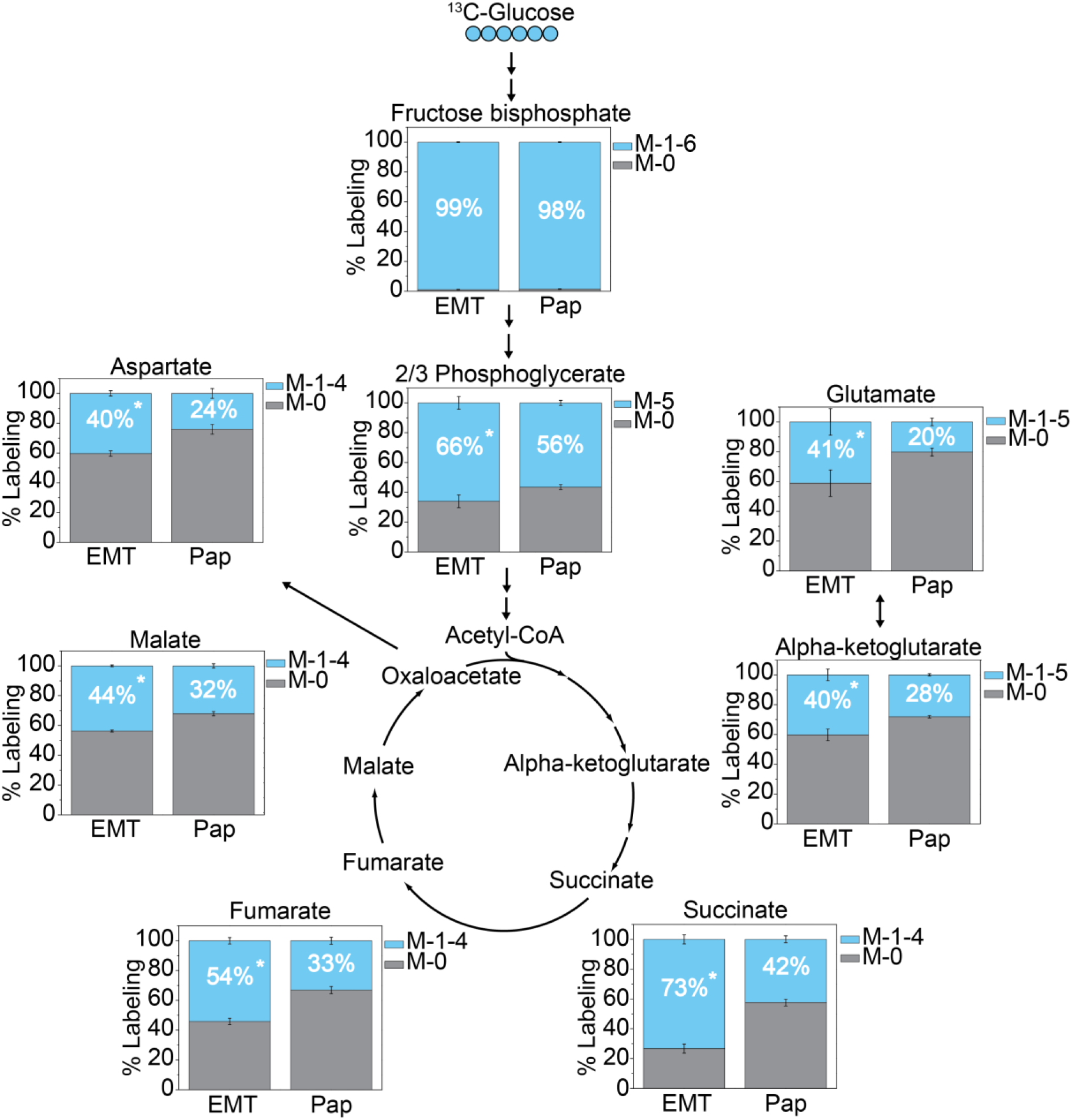
^13^C-Isotope labeling from glucose into the TCA cycle is significantly higher in the EMT subtype. Cells were incubated in ^13^C-glucose containing media for 4 h and extracted for metabolites. Grey boxes represent the unlabeled (M-0 isotopologue) proportion for each metabolite. Blue boxes represent the sum of all potential labeled isotopologues (M-1 through M-*X*, where *X* represents the total number of carbons in the metabolite) for each metabolite. Data are displayed as means ± S.D., N = 3 (*p-value < 0.05).

### Isotope labeling into nucleotide biosynthesis is elevated in the papillary subtype

Compared to the EMT cells, the papillary cells exhibit increased levels of nucleotides ADP and ATP, as well as ribulose-5-phosphate and ribose-5-phosphate, two intermediates in the PPP (**Figure 2C-D**). To determine whether these measurements reflect increased nucleotide production or decreased nucleotide consumption, we applied the same isotope labeling techniques described above. Nucleotides can be generated through de novo biosynthesis or salvage pathways. Several carbon sources contribute to the formation of purine and pyrimidine rings during *de novo* biosynthesis. Purine carbons are derived from glycine (2 carbons), formate (2 carbons), and bicarbonate (1 carbon). Pyrimidine carbons are derived from aspartate (3 carbons, predominately from glutamine metabolism) and bicarbonate (1 carbon).^22^ Salvage pathways recycle intermediates scavenged from the environment or produced from RNA and DNA degradation to generate nucleotides, and these pathways require less energy per produced nucleotide compared to *de novo* biosynthesis. The carbon sources for purine and pyrimidine nucleotides are highlighted in **Figure 4A,B**.

Isotope labeling studies show that the papillary cells have higher *de novo* nucleotide biosynthesis compared to the EMT cells (**Figure 4C,D; Supplementary Tables 1-2**). When cells are fed ^13^C-glucose, the M-5 isotopologue of inosine monophosphate (IMP) and ATP can be derived from either *de novo* or salvage pathways, while all other isotopologues of IMP and ATP (M-1 to M-4 and M-6 to M-10, referred to as M-Other in **Figure 4C**) can only be derived through *de novo* biosynthetic pathways (**Figure 4A**). As shown in **Figure 4C** and **Supplementary Table 1**, M-Other is higher in the papillary cells for both IMP (23% in papillary vs. 19% in EMT) and ATP (19% in papillary vs. 15% in EMT). Further, ^13^C-glutamine labeling shows increased levels of the M3 isotopologue of uridine triphosphate (UTP) in the papillary cells (23% in papillary vs. 17% in EMT; **Figure 4D; Supplementary Table 2**) – this isotopologue can also only be derived from *de novo* biosynthesis (**Figure 4B**). Therefore, the papillary cells demonstrate increased *de novo* biosynthesis of both purine and pyrimidine nucleotides compared to the EMT cells. Notably, we find no difference between EMT and papillary cells in ^13^C-glucose labeling into ribose-5-phosphate, serine, and glycine as well as ^13^C-glutamine labeling into aspartate (**Supplementary Figure 2**), which indicates that increased nucleotide biosynthesis in the papillary cells is not simply due to greater abundance of labeled precursors for these pathways. Increased *de novo* nucleotide biosynthesis could reflect a preference to utilize this metabolic pathway to generate nucleotides in the papillary subtype. This would indicate targeting *de novo* nucleotide biosynthesis is likely to be more effective in the papillary subtype.

**Figure 4:**
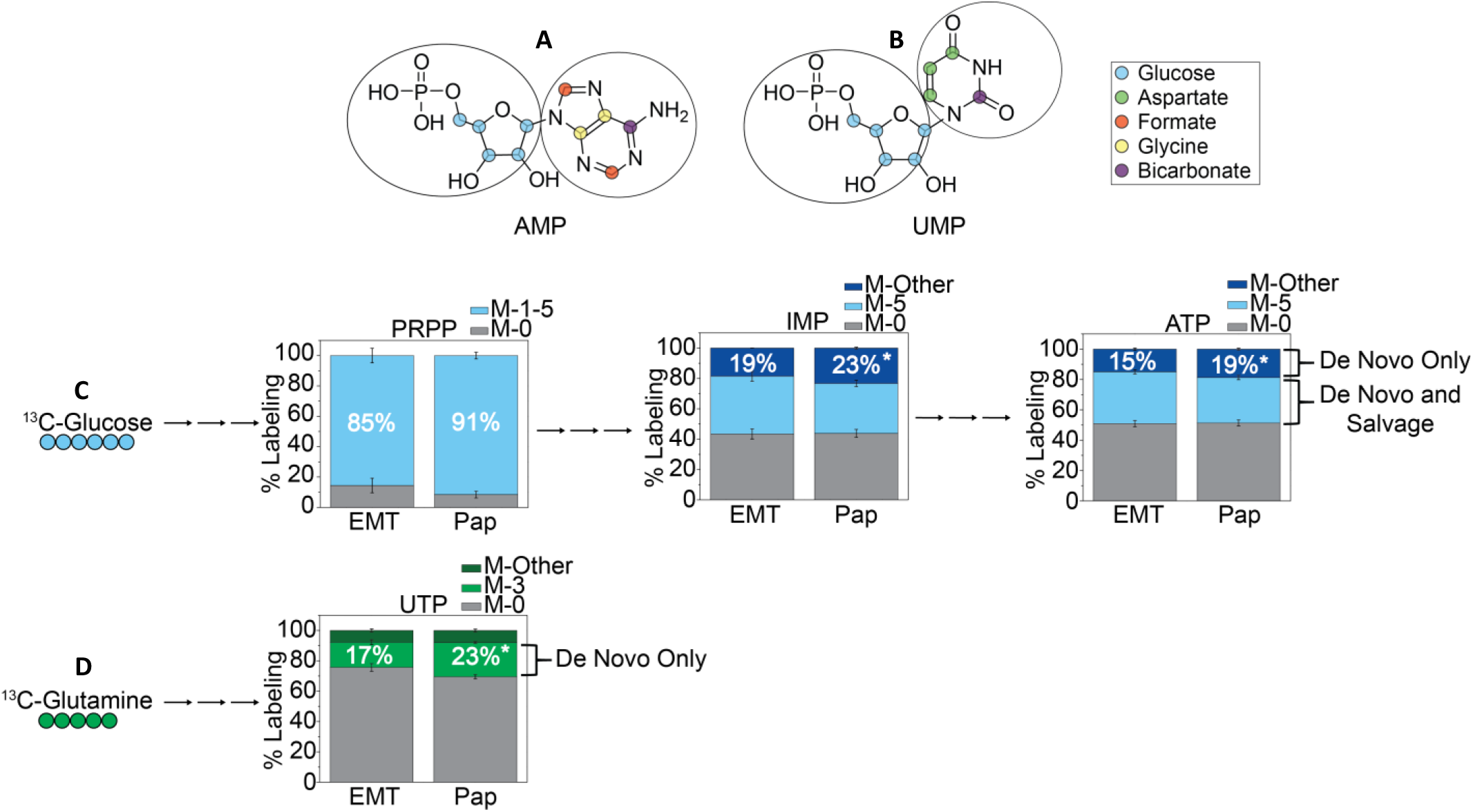
^13^C-Isotope incorporation from glucose and glutamine into nucleotide biosynthesis is higher in the papillary subtype. Molecular diagrams of (**A**) purine nucleotide AMP and (**B**) pyrimidine nucleotide UMP with carbon sources highlighted as colored circles. The 5 carbon ribose sugar of both (**A,B** left circle) is derived from glucose metabolism. Isotopologues of this mass (M-5) reflect both salvaged nucleotides and nucleotides produced by *de novo* biosynthetic pathways when ^13^C-glucose is administered. The 5 carbons comprising the purine ring of AMP (**A**, right circle) are derived from glycine, formate, and bicarbonate, all three of which can also be derived from glucose metabolism. Therefore, when ^13^C-glucose is administered isotopologues of other masses (M-1 to M-4 and M-6 to M-10, referred to as M-Other) reflect only purine production by *de novo* biosynthesis. The 4 carbon comprising the pyrimidine ring of UMP (**B**, right circle) are derived from bicarbonate and aspartate. Aspartate is predominantly derived from glutamine metabolism and provides 3 carbons to UMP; therefore, when ^13^C-glutamine is administered, M-3 isotopologues reflect de novo UMP biosynthesis. (**C**) ^13^C-Glucose labeling into PRPP, IMP, and ATP. (**D**) ^13^C-Glutamine labeling into UTP. Cells were incubated in ^13^C-glucose (blue) or ^13^C-glutamine (green) containing media for 4 h and extracted for metabolites. Grey boxes represent the unlabeled proportion for each metabolite at 4 h. Colored boxes represent isotopologues for each metabolite and are sorted based on carbon source. Data are displayed as means ± S.D., N = 3 (*p-value < 0.05).

### Relative metabolic pathway activity correlates with drug response

To test whether metabolism-targeting drugs impact cell proliferation in a subtype-specific manner, cell proliferation was determined in the presence of metabolism-targeting compounds. Compounds were chosen based on our initial findings that the EMT subtype increased glutathione biosynthesis and TCA cycle metabolism, while the papillary subtype increased *de novo* nucleotide biosynthesis. The three selected compounds were: 1) Buthionine sulfoximine (BSO), an inhibitor of gamma-glutamylcysteine synthetase in glutathione biosynthesis;^25^ 2) CPI-613, which targets pyruvate dehydrogenase and alpha-ketoglutarate dehydrogenase in the TCA cycle;^26,27^ and 3) 5-Fluorouracil (5FU) which is an inhibitor of thymidylate synthase in *de novo* nucleotide biosynthesis (**Supplementary Figure 3**).^28,29^ We found that targeting each distinct metabolic feature inhibits breast cancer cell proliferation in a subtype-specific manner. Since the EMT cells display increased levels of both oxidized and reduced glutathione compared to papillary cells (**Figure 2B**), they are more likely to be sensitive to glutathione biosynthesis inhibition. As expected, targeting glutathione biosynthesis with BSO was more effective at inhibiting proliferation of the EMT cells vs. the papillary cells (**Figure 5A-B**). Consistently, the IC_50_ for this compound was significantly lower for the EMT cells (34 μM) vs. the papillary cells (49 μM; p-value <0.0001; **Supplementary Figure 4A,D**). The EMT cells also have higher TCA cycle activity compared to the papillary cells (**Figure 3**) and should therefore be more sensitive to TCA cycle inhibition. Indeed, targeting TCA cycle metabolism with CPI-613 was more effective in the EMT vs. papillary cells (**Figure 5C-D**); the IC_50_ for this compound was also significantly lower for the EMT (123 μM) vs. the papillary cells (153 μM; p-value <0.0001; **Supplementary Figure 4B,D**). Finally, the papillary subtype demonstrates increased *de novo* nucleotide biosynthesis (**Figure 4**) and should therefore be most sensitive to compounds which target nucleotide biosynthesis. Indeed, we find that targeting nucleotide metabolism with 5FU was most effective at inhibiting proliferation of the papillary cells vs. the EMT cells (**Figure 5E-F**); the IC_50_ for this compound was significantly lower for the papillary cells (397 nM) vs. the EMT cells (1359 nM; p-value <0.0001; **Supplementary Figure 4C-D**). These results illustrate how metabolic profiles can be used to identify therapeutic targets for subtypes of breast cancer.

**Figure 5:**
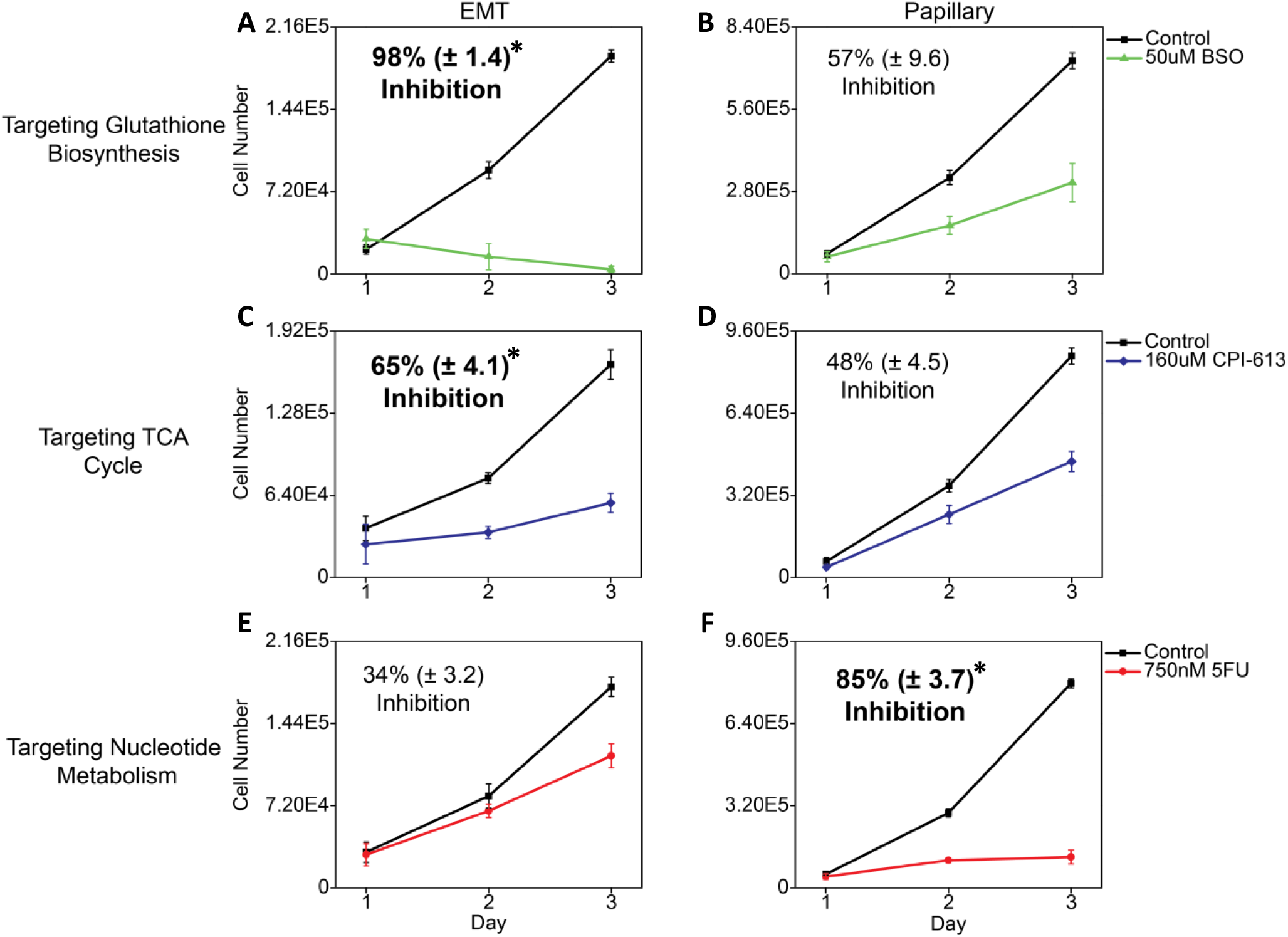
Metabolism targeting drugs have subtype-specific effects on cell proliferation. Cells from each subtype were seeded into vehicle or drug containing media, and viable cells were counted daily for 3 days. Proliferation inhibition is determined using the ratio of the drug treated cell count at day 3 to the vehicle treated cell count at day 3. Bolded values indicate the subtype most affected by each compound. Data are displayed as means ± S.D., N = 3 (*p-value < 0.01).

## DISCUSSION

In this study, we demonstrate the utility of targeting subtype-specific metabolic profiles to inhibit cancer cell proliferation. Using a combination of unlabeled and isotope-labeled massspectrometry-based metabolomics techniques, we developed comprehensive metabolic profiles of two histologically distinct breast cancer subtypes derived from the MMTV-Myc mouse model. We further leveraged these metabolic profiles to identify therapeutic targets for each subtype, and demonstrate that inhibiting cancer cell metabolism is most effective when tailored to the underlying metabolic profile of the cancer in question. This approach, when translated to human disease, has the potential to improve patient outcomes, as it will lead to development of novel metabolic drugs for cancer subtypes that currently lack targeted therapies. This may be particularly relevant for the EMT subtype, as it has been correlated with the claudin-low subtype of breast cancer in humans,^15^ a subtype which generally carries a poor prognosis and currently lacks targeted therapeutic options.^16^

In recent years, there has been growing interest in taking advantage of altered metabolism in cancer for treatment.^30–32^ Our findings underscore the importance of metabolic profiling as a tool for designing novel personalized therapies. Of the three compounds evaluated in this study, 5FU is currently approved as a chemotherapeutic agent, and is used to treat a variety of malignancies.^29^ BSO has demonstrated utility as a sensitizing agent in pre-clinical models of anti-endocrine therapy resistant ER+ breast cancer,^33^ and in multiple myeloma treated concurrently with the chemotherapeutic melphalan.^34^ More recently, BSO has shown promise in early clinical trials as a chemosensitizing agent in combination with melphalan for treatment of pediatric neuroblastoma.^35,36^ Finally, CPI-613 is being investigated as a component in combination therapies for several malignancies including colorectal cancer,^37^ small cell lung cancer,^38^ and pancreatic cancer.^39^ Our findings support investigating these compounds to treat specific subtypes of breast cancer, as each tested compound demonstrates some degree of inhibition regardless of subtype. Moreover, our findings provide an additional rationale for subtype-specific drug selection based on the underlying metabolism of the cancer cells in question, as the drug sensitivity for the EMT and papillary cells directly correlates with the metabolic profile of each subtype.

Our results may also in part explain why some cancer patients do not respond to a given metabolism-targeting therapy. For example, 5FU as a monotherapy to treat metastatic colorectal cancer demonstrates a response rate of only 10-20%,^40^ indicating a significant proportion of patients fail to respond to 5FU therapy. Response rates are better when 5FU is used in combination therapies to treat metastatic breast cancer, with response rates of 40-80% depending on the specific combination therapy.^40^ In such cases, it is possible the metabolic profile of the poor responder’s cancer differs significantly from the metabolic profile of a cancer that responds well to treatment. Our study demonstrates that 5FU will be most effective in cancer subtypes that upregulate *de novo* nucleotide biosynthesis. This may be particularly relevant for cancers with elevated Myc activity, since Myc regulates the expression of numerous genes in nucleotide biosynthesis including thymidylate synthase (TYMS),^22^ the primary target of 5FU. Increased Myc expression has been observed in TNBC compared to hormone receptor positive breast cancer,^41^ and patients with TNBC have better response rates to neoadjuvant chemotherapy regimens that contain 5FU.^42^ Further, Myc overexpression in hepatocellular carcinoma has recently been shown to decrease both oxidized and reduced glutathione levels in tumor tissue by downregulating glutathione biosynthesis genes.^43^ Our findings support this in breast cancer, as the papillary cells, which have relatively higher Myc activity, demonstrate decreased glutathione levels compared to EMT cells.^14^ This is consistent with papillary cells being less sensitive to BSO treatment. Thus, breast cancers with increased Myc expression may be less likely to respond to therapies that target glutathione biosynthesis, and breast cancers that lack Myc overexpression may respond favorably to glutathione biosynthesis inhibitors. Other metabolic features associated with increased Myc signaling, such as increased glutaminolysis^11,44^ and fatty acid metabolism,^45^ are also under investigation as potential therapeutic targets. Therefore, metabolomic analysis of patient samples could provide clinicians with additional prognostic information to guide treatment plans, ultimately improving patient outcomes while decreasing unnecessary side effects by avoiding ineffective treatment regimens.

## METHODS

### Primary mouse tumors

All animal use was performed in accordance with institutional and federal guidelines. Primary MMTV-Myc EMT and MMTV-Myc papillary tumors were acquired as a gift from Dr. Eran Andrechek and have been previously described.^14^ Tumors were sectioned, formalin-fixed, and paraffin embedded for histological examination with hematoxylin and eosin staining. Tumor derived cell lines were established by mechanical dissociation of primary tumors using scissors, followed by culturing tumor pieces in cell culture media.^46^

### Cell lines and culture conditions

EMT and papillary tumor derived cell lines were cultured in Dulbecco’s Modified Eagle Medium (DMEM Corning, Corning, New York 10-017-CM) with 25 mM glucose without sodium pyruvate supplemented with 2 mM glutamine (Corning, 25-005-CI) 10% heat-inactivated fetal bovine serum (MilliporeSigma, Burlington Massachusetts, 12306C), and 1% penicillin and streptomycin (Corning, 30-002-CI). Cells were maintained at 37°C with 5% CO_2_.

### Metabolic profiling

Unlabeled, targeted metabolomics was performed as previously described.^23^ Briefly, cells were seeded in 6-well tissue culture plates at 50,000 cells/well and cultured for 48 hours. Cells were washed with saline (VWR, Radnor, Pennsylvania, 16005-092) and metabolism was quenched with addition of cold methanol. The final metabolite extraction solvent ratios were methanol:water:chloroform (5:2:5). The polar phase was collected and dried under a stream of nitrogen gas. The dried metabolites were then resuspended in HPLC-grade water for analysis. LC-MS analysis was performed with ion-pairing reverse phase chromatography using an Ascentis Express column (C18, 5 cm × 2.1 mm, 2.7 μm, MilliporeSigma, 53822-U) and a Waters Xevo TQ-S triple quadrupole mass spectrometer. Mass spectra were acquired using negative mode electrospray ionization operating in multiple reaction monitoring (MRM) mode. Peak processing was performed using MAVEN^47^ and data for each sample was normalized to the median signal intensity. Heatmaps were generated using Cluster 3.0^48^ and exported using Java Treeview.^49^

### Isotope labeling studies

For isotope labeling experiments, DMEM without glucose or glutamine was prepared from powder (MilliporeSigma, D5030) and supplemented with either ^13^C-glucose (Cambridge Isotope Laboratories, Tewksbury, Massachusetts, CLM-1396) and unlabeled glutamine (MilliporeSigma, G8540) or unlabeled glucose (Fisher Scientific, Hampton, New Hampshire, D16) and ^13^C-glutamine (Cambridge Isotope Laboratories, CLM-1822). Cells were then seeded and cultured as described above. Prior to metabolite extraction, media was switched to isotope containing media and samples were collected at T = 0 (unlabeled) and 240 minutes. Metabolite extraction and analysis were performed as above. Labeling data was corrected for natural isotope abundance using IsoCor.^50^

### Cell proliferation and drug response studies

Cells were seeded at a density of 20,000 cells/well in 12-well tissue culture plates and treated with either vehicle (DMSO, MilliporeSigma, D4540) or the indicated drugs. CPI-613 (Cayman Chemical, Ann Arbor, Michigan, 16981), buthionine sulfoximine (Cayman Chemical, 14484), and 5-fluorouracil (TCI, Tokyo, Japan, F0151). Cells were counted daily for 3 days using a Nexcelom Cellometer Auto T4 cell counter and viable cells were determined using trypan blue exclusion (VWR, 45000-717). Proliferation inhibition was determined using the ratio of the drug treated cell count at day 3 to the vehicle treated cell count at day 3.

### Statistical analyses

Statistical analyses were performed using unpaired Student’s t-test except where otherwise noted. P-values were adjusted in R using the p.adjust() function to account for multiple testing using the Benjamini-Hochberg procedure. All error bars presented are standard deviation. IC_50_ values and statistical analysis of drug response were calculated using nonlinear regression performed by GraphPad Prism.

## ACKNOWLEDGEMENTS

The authors thank Deanna Broadwater, Elliot Ensink, Hyllana Medeiros, Shao Thing Teoh, and Lei Yu for helpful discussions and critical reading of this manuscript. The authors thank Eran Andrechek for the generous gift of primary MMTV-Myc EMT and MMTV-Myc papillary tumors. The authors also thank the MSU Mass Spectrometry and Metabolomics Core and the MSU Investigative HistoPathology Laboratory. **Funding:** This work was supported by the Office of the Assistant Secretary of Defense for Health Affairs, through the Breast Cancer Research Program, under Award No. W81XWH-15-1-0453.

## COMPETING INTERESTS

The authors declare that they have no competing interests.

## Supplementary figures and tables

**Supplementary Figure 1:**
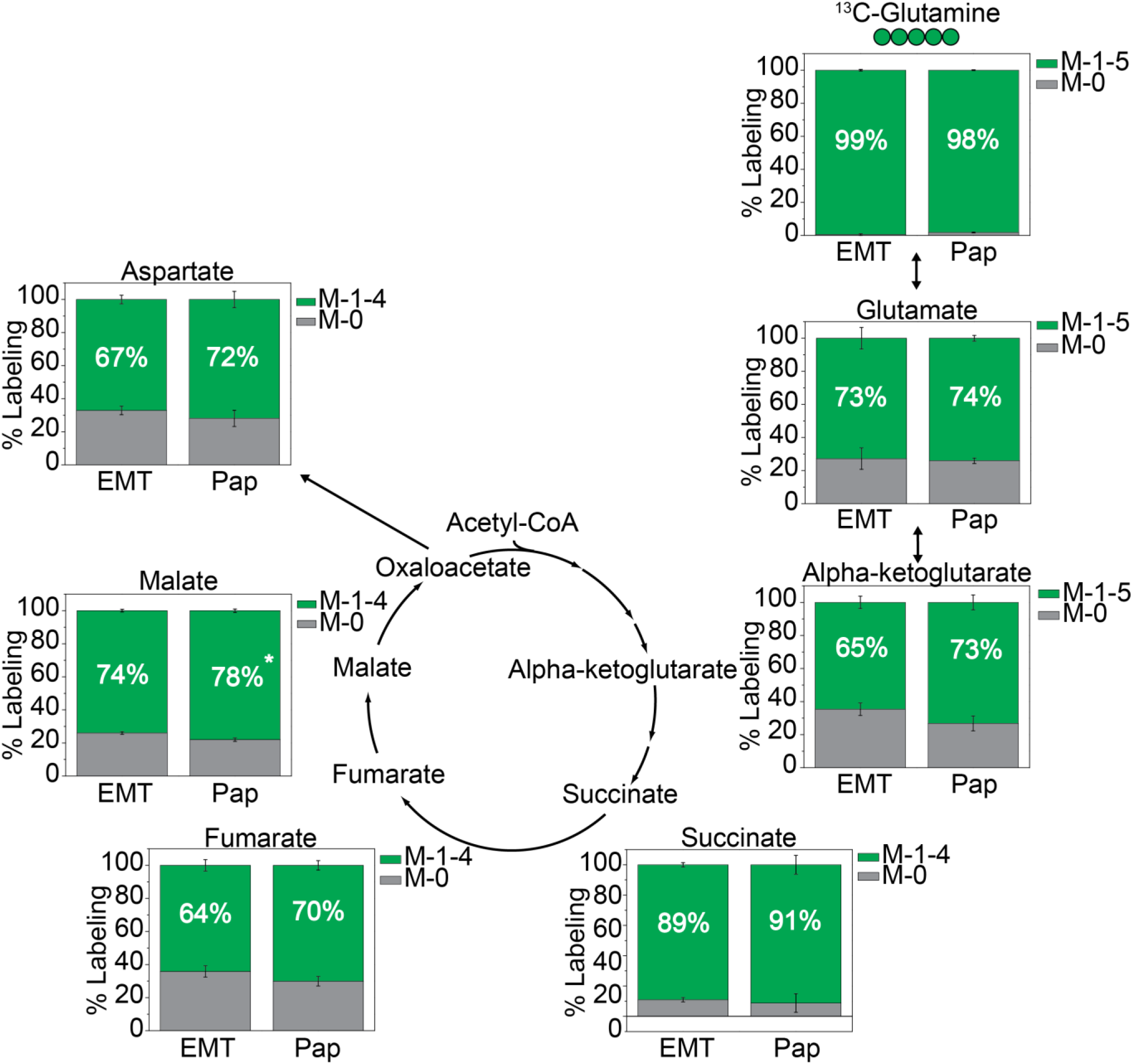
^13^C-Isotope labeling from glutamine into the TCA cycle is similar between subtypes. Cells were incubated in ^13^C-glutamine containing media for four hours and extracted for metabolites. Grey boxes represent the unlabeled proportion for each metabolite at four hours. Green boxes represent the sum of all potential isotopologues for each metabolite. Data are displayed as means ± S.D., N = 3 (Statistically significant differences (p-value < 0.05) are marked with asterisks (*).

**Supplementary Figure 2:**
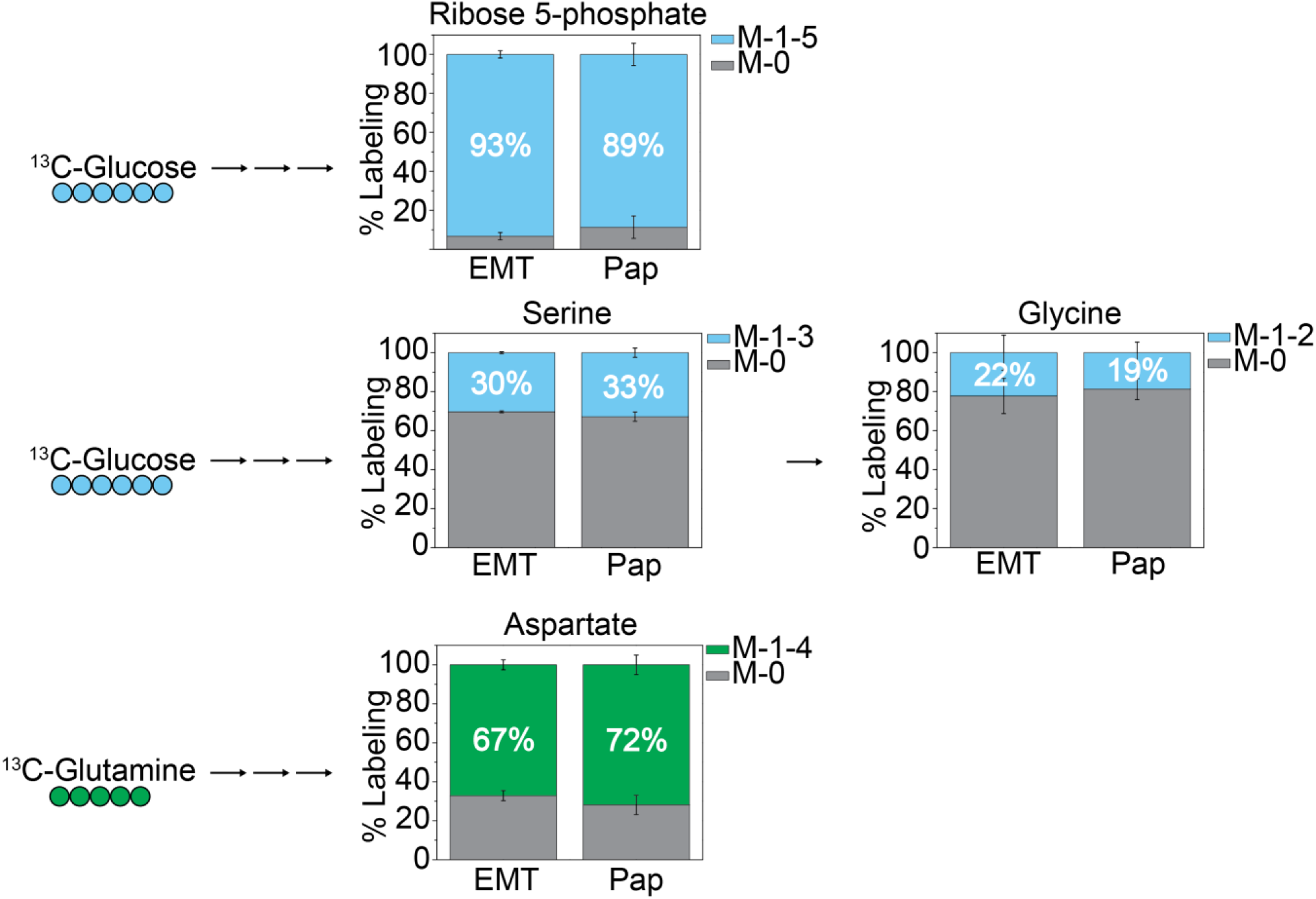
^13^C-Isotope labeling from glucose into ribose 5-phospahte, serine, and glycine and from glutamine into aspartate is similar between subtypes. Cells were incubated in ^13^C-glucose (blue) or ^13^C-glutamine (green) containing media for 4 h and extracted for metabolites. Grey boxes represent the unlabeled proportion for each metabolite at four hours. Colored boxes represent isotopologues for each metabolite and are sorted based on carbon source. Data are displayed as means ± S.D., N = 3 (Statistically significant differences (p-value < 0.05) are marked with asterisks (*).

**Supplementary Figure 3:**
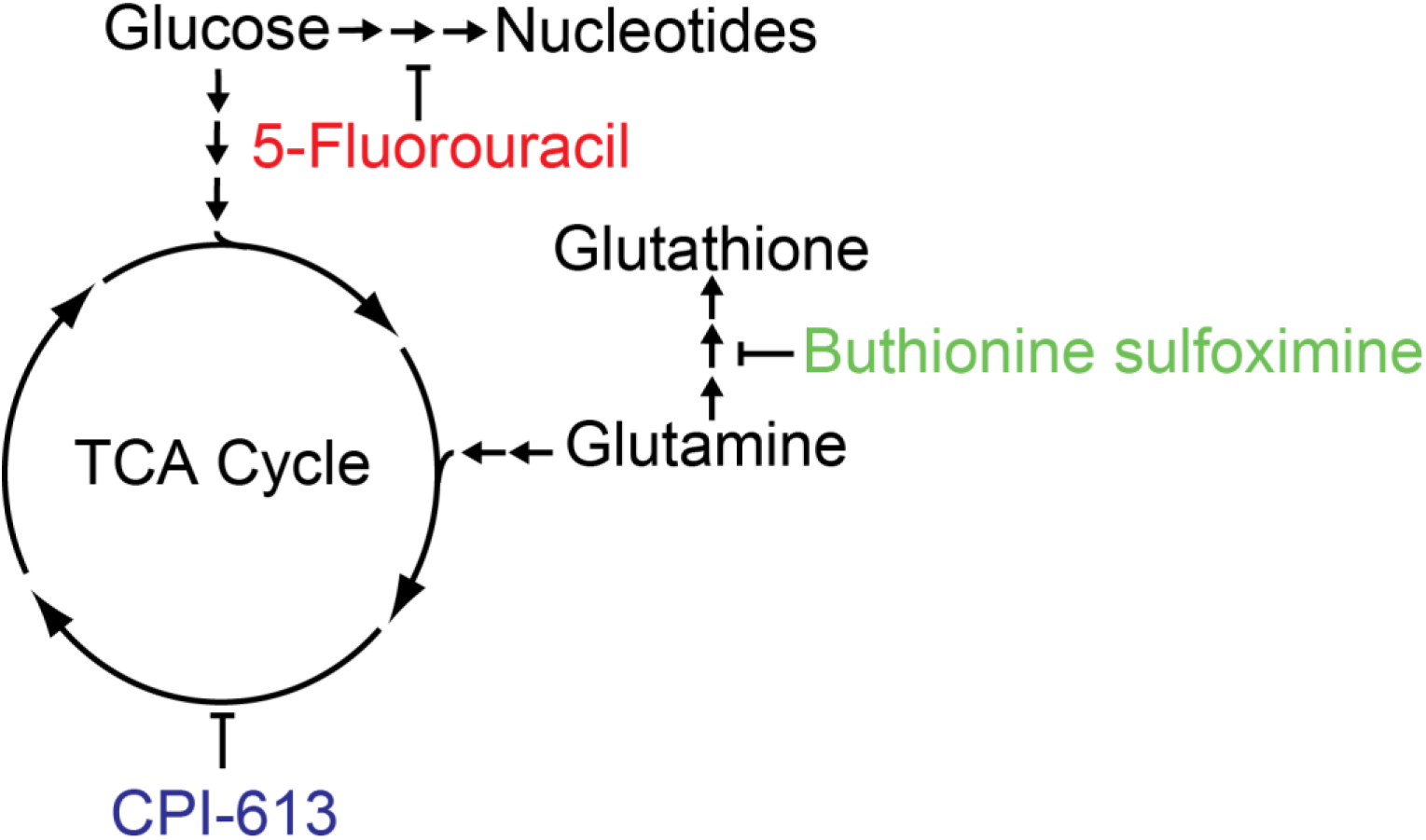
Schematic overview of metabolism targeting drugs and affected pathways. Compounds were chosen based on the metabolic differences identified in the EMT and papillary subtypes. Buthionine sulfoximine targets gamma-glutamylcysteine synthetase in glutathione biosynthesis, CPI-613 targets pyruvate dehydrogenase and alpha-ketoglutarate dehydrogenase in the TCA cycle, and 5-fluorouracil targets thymidylate synthase in nucleotide biosynthesis.

**Supplementary Figure 4:**
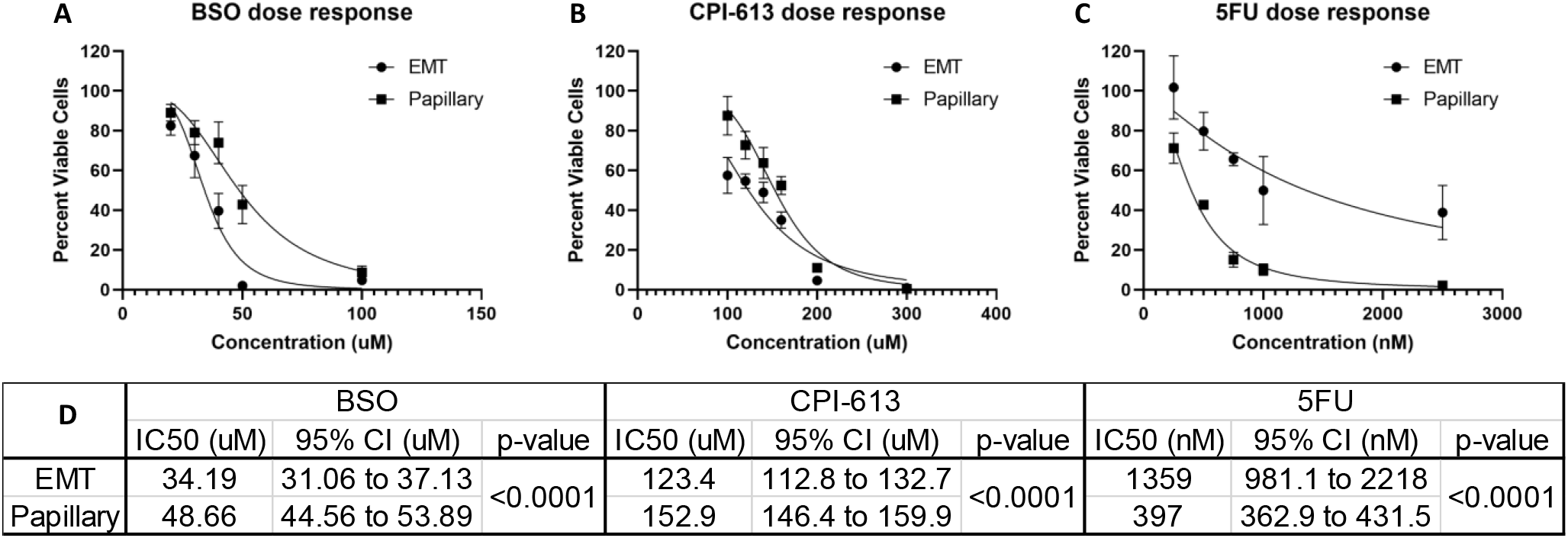
Dose response curves for metabolism targeting drugs. (A) Buthionine sulfoximine (BSO) and (B) CPI-613 demonstrate greater effects on the EMT subtype. (C) 5-fluorouracil (5-FU) demonstrates a greater effect on the papillary subtype. (D) IC_50_ and 95% confidence intervals calculated from each dose response curve. Nonlinear regression and statistical analyses performed by GraphPad Prism. Data are means ± S.D. (n=3).

**Supplementary Table 1:**
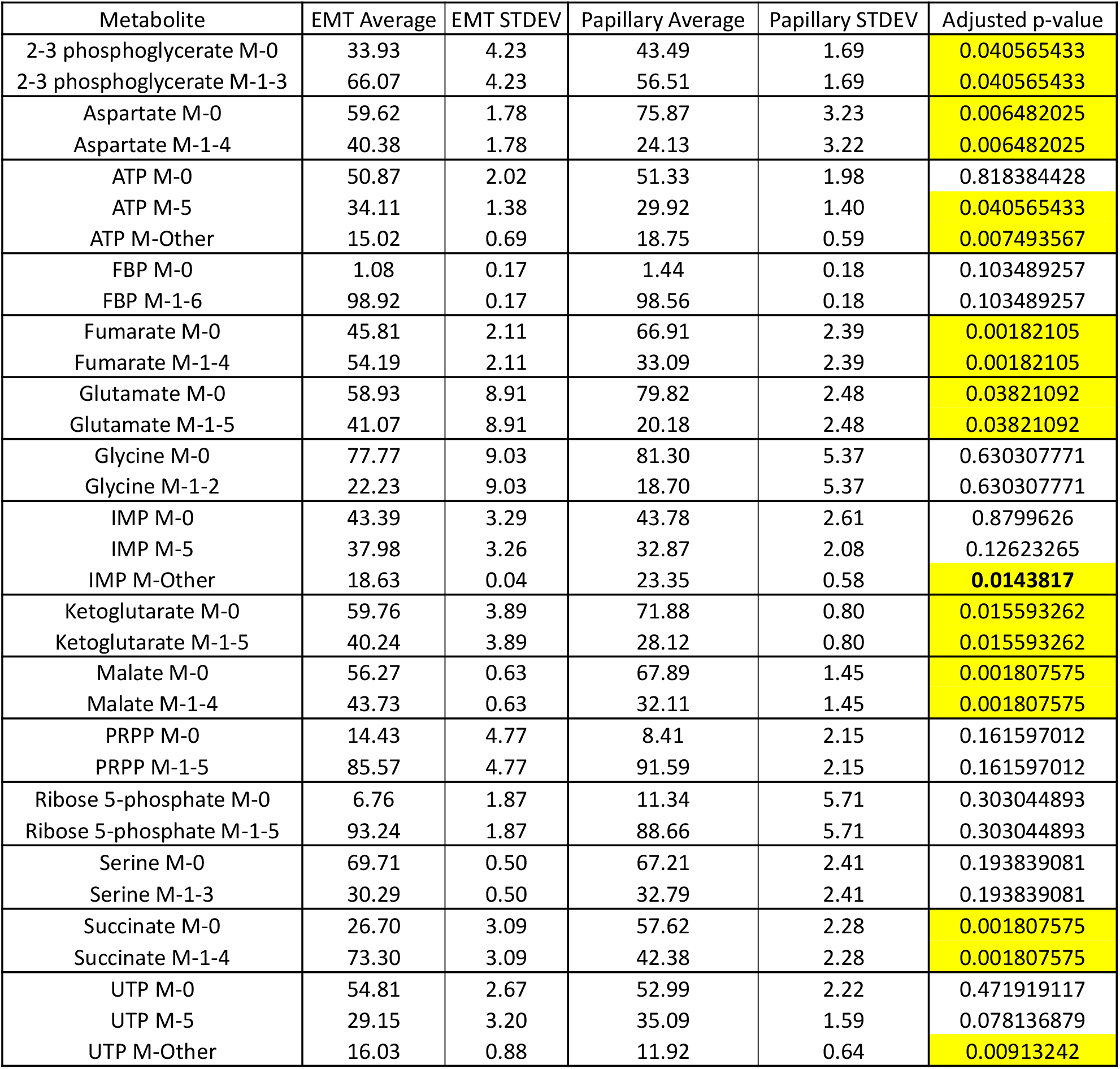
^13^C-Isotope percent labeling from glucose with statistical significance. Data represent means and S.D. of 3 replicates. Unpaired Student’s t-tests were used to determine p-values. Bold values indicate Welch’s t-test was used. P-values were adjusted to account for multiple testing using the Benjamini-Hochberg procedure. Highlighted values are statistically significant with adjusted p-value < 0.05.

**Supplementary Table 2:** ^13^C-Isotope percent labeling from glutamine with statistical significance. Data represent means and S.D. of 3 replicates. Unpaired Student’s t-tests were used to determine p-values. P-values were adjusted to account for multiple testing using the Benjamini-Hochberg procedure. Highlighted values are statistically significant with adjusted p-value < 0.05.

| Metabolite | EMT Average | EMT STDEV | Papillary Average | Papillary STDEV | Adjusted p-value |
| --- | --- | --- | --- | --- | --- |
| Aspartate M-0 | 32.87 | 2.54 | 28.11 | 4.91 | 0.29728481 |
| Aspartate M-1-4 | 67.13 | 2.54 | 71.89 | 4.91 | 0.29728481 |
| Fumarate M-0 | 35.84 | 3.42 | 29.89 | 2.84 | 0.13841247 |
| Fumarate M-1-4 | 64.16 | 3.42 | 70.11 | 2.84 | 0.13841247 |
| Glutamate M-0 | 27.18 | 6.49 | 25.91 | 1.67 | 0.7711491 |
| Glutamate M-1-5 | 72.82 | 6.49 | 74.09 | 1.67 | 0.7711491 |
| Glutamine M-0 | 0.55 | 0.48 | 1.82 | 0.25 | 0.05282444 |
| Glutamine M-1-5 | 99.45 | 0.48 | 98.18 | 0.25 | 0.05282444 |
| Ketoglutarate M-0 | 35.37 | 3.76 | 26.79 | 4.46 | 0.13531873 |
| Ketoglutarate M-1-5 | 64.63 | 3.76 | 73.21 | 4.46 | 0.13531873 |
| Malate M-0 | 26.03 | 0.82 | 22.00 | 1.02 | 0.0335597 |
| Malate M-1-4 | 73.97 | 0.82 | 78.00 | 1.02 | 0.0335597 |
| Succinate M-0 | 10.95 | 1.40 | 8.78 | 6.18 | 0.70940041 |
| Succinate M-1-4 | 89.05 | 1.40 | 91.22 | 6.18 | 0.70940041 |
| UTP M-0 | 75.60 | 2.53 | 69.45 | 1.37 | 0.05883558 |
| UTP M-3 | 16.68 | 1.55 | 22.60 | 0.51 | 0.0335597 |
| UTP M-Other | 7.72 | 0.99 | 7.95 | 0.86 | 0.7711491 |

